# GENLIB: new function to simulate haplotype transmission in large complex genealogies

**DOI:** 10.1101/2022.10.28.514245

**Authors:** Mohan Rakesh, Hélène Vézina, Catherine Laprise, Ellen E Freeman, Kelly M Burkett, Marie-Hélène Roy-Gagnon

## Abstract

**Summary:** Founder populations with deep genealogical data are well suited for investigating genetic variants contributing to diseases. Here, we present a new function added to the genealogical analysis R package GENLIB, which can simulate the transmission of haplotypes from founders to probands along very large and complex user-specified genealogies.

**Availability and implementation:** The new function is available in the latest version of the GENLIB package (v1.1.6), available on the CRAN repository and from https://github.com/R-GENLIB/GENLIB. Stand-alone scripts for analyzing the output of the function can be accessed at https://github.com/R-GENLIB/simuhaplo_scripts.

## Introduction

Founder populations have been utilized extensively in the study of Mendelian diseases because they can have higher incidence rates of rare autosomal-recessive genetic diseases due to drift effects, e.g.: Gaucher disease, Tay-Sachs disease, and cystic fibrosis in the Ashkenazi Jewish population (Charrow, 2004), or one of the over 30 identified autosomal recessive diseases with elevated frequency in the Finnish population (Norio, 2003; Pastinen, et al., 2001). In founder populations, affected individuals are more likely to have the causal mutation on longer haplotypes that are homozygous by recent decent, aiding in mutation discovery (Bourgain and Genin, 2005; Libiger and Schork, 2007). Some founder populations have extensive records allowing for reconstruction of deep and large genealogies (Falchi, et al., 2004; Liu, et al., 2007; Ober, et al., 2001; Vézina and Bournival, 2020). Gene-dropping simulations (Chen, et al., 2015; Maccluer, et al., 1986) can be performed within these genealogies, wherein ancestral genotypes are passed down a fixed pedigree structure. For example, allele-dropping was used to study mutation frequencies in the Hutterite (Chong, et al., 2012), and French-Canadian (Heyer, 1999) founder populations.

Gene-dropping is not limited to dropping specific alleles. Transmission of genomic regions, chromosomes, or even the entire genome can be simulated. This type of simulations can provide important information on the distribution of genomic sharing and the probability of sharing a specific genomic segment among close or distant relatives, and can identify specific founders and transmission paths responsible for the observed sharing. However, in very large genealogies, these gene-dropping simulations are computationally feasible only if one does not consider the allelic state of any specific locus, but rather only the positions of recombination events and the origin (founder) of the segments bounded by the crossovers (Cheng, et al., 2015). We have implemented such a gene-dropping simulation tool in the GENLIB genealogical analysis R package (Gauvin, et al., 2015; R Core Team, 2021). This new tool (named gen.simuhaplo) is fast even for large genomic regions and deep genealogies with many individuals because it does not consider any alleles, mutations, or phenotypes. To our knowledge, it is the first user-friendly simulation tool that can perform gene-dropping simulations of long genomic segments in very large and complex genealogies, such as those available in the French-Canadian founder population, while allowing the ability to retrace all transmission paths.

### Implementation and usage

The gen.simuhaplo function is written in C++ and uses the Rcpp package (Eddelbuettel and Francois, 2011), as well as the R core C API to interface with the R environment.

The function simulates meiosis using one of three possible models (see Supplementary Appendix 1 for details). The first is a Poisson process, which is a no-interference model of meiosis (Haldane, 1919). The second model is a count-location model (Karlin and Liberman, 1978; Karlin and Liberman, 1979), where the number of chiasma is sampled from a zero-truncated Poisson distribution (Risch and Lange, 1979; Sturt, 1976). This model accounts for the obligate chiasma phenomenon, i.e., requirement for at least one chiasma per tetrad (Fledel-Alon, et al., 2009). The third model is a stationary gamma process (Broman and Weber, 2000), which accounts for chromosomal interference. After the locations of the crossovers are obtained in Morgans they are converted from genetic distance to physical distance and a meiotic product is selected and transmitted. The user may provide a map to convert genetic distance to physical distance, or else the relationship between genetic and physical distance will be assumed to be linear across the length of the chromosome. The choice of model and the use of a genetic-physical map can alter the distribution of the lengths of segment identical-by-descent (IBD) (Caballero, et al., 2019).

After importing the genealogical data and creating a “genealogy” object within GENLIB, the gen.simuhaplo function can be called. First, a list of individuals is created such that any dependent individual is listed after their ancestors (i.e.: parents before children, and all ancestors before descendants). Since a family tree is a directed acyclic graph, we can always perform such a topological sort. Then we iterate through the list, and, at each individual, we simulate meiosis in the parents and pass down the meiotic products. A meiotic product is created by first obtaining a list of chiasma positions (in base pairs, BP), then assuming that each chiasma will have a probability of 0.5 of appearing in the given meiotic product as a crossover. After selecting all the crossovers, we select the parental copy with which the meiotic product begins with probability of 0.5. We copy over the parental chromosome until reaching a crossover position, then we alternate to copying from the other parental chromosome. The chromosomes are stored as linked lists where each segment contains its end position in BP, and points to the next segment in the chromosome.

The output of the function is a text file containing the location of the breakpoints and the code for the founder haplotypes, from which the simulated haplotypes can be derived for all the specified probands. Optionally the function can output a second text file containing the haplotypes for all individuals along the inheritance paths. The optional second text file contains a recombination history that allows users to easily retrace genomic segments from probands up to internal ancestors (see Supplementary Appendix 2). An example of the output format is shown in Figure 1. Length of segments shared, or position shared on the simulated segment, can be obtained from this output even though no actual genotypes at specific markers in the segment are used in the simulations. If users need to simulate with genotype data, we provide a Perl script for converting the output into genotype data (see Supplementary Appendix 2). The user can provide haploid genotypes for the founders and the proband haplotypes can be converted into corresponding genotype data (phasing into haplotypes can then be ignored if desired). This could be used to distinguish between alleles shared identical by descent versus by state.

**Figure 1.**
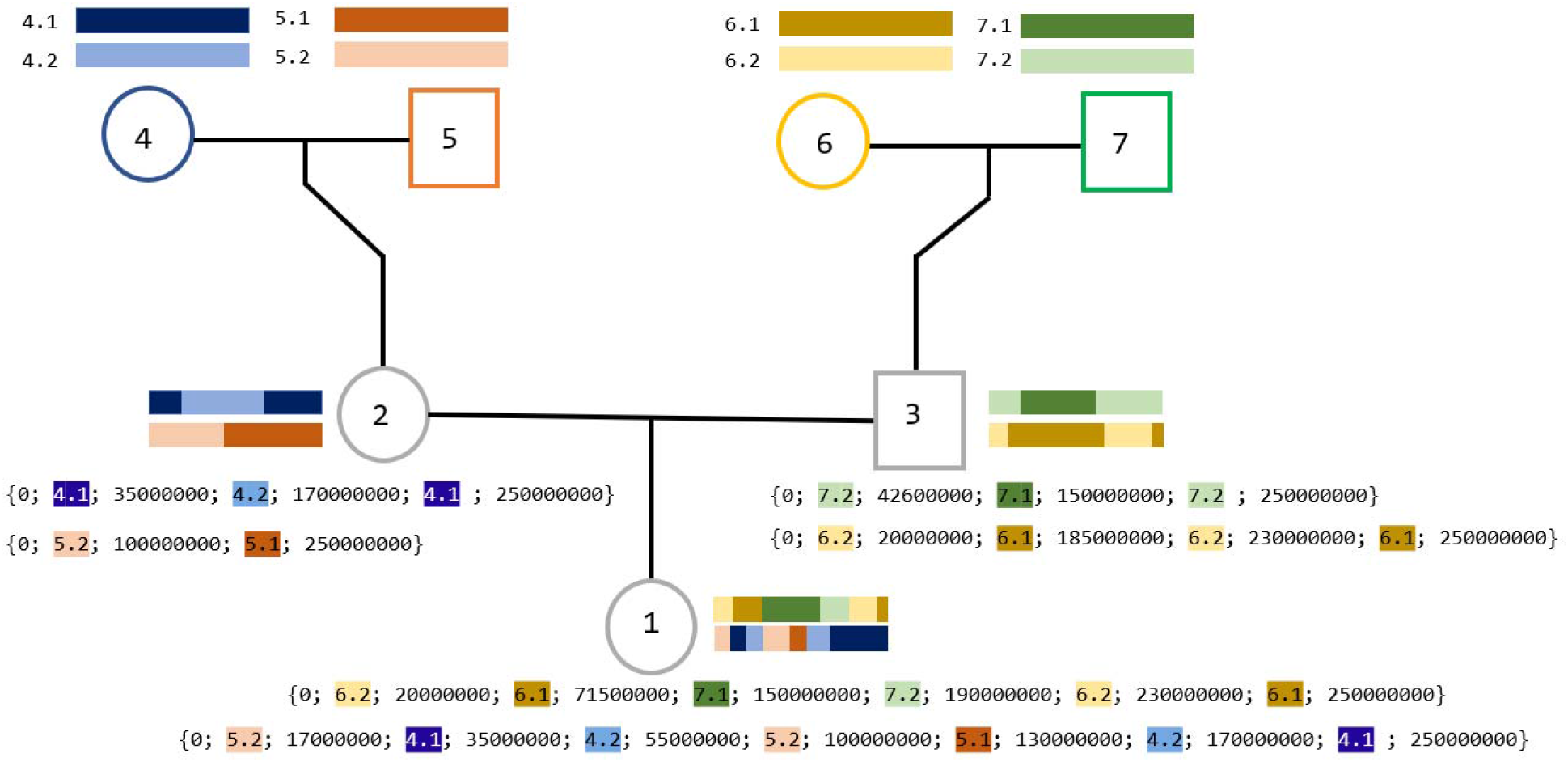
Example simulation of a hypothetical 250,000,000 BP segment. Each individual in the genealogy has a unique integer ID. This ID number is used to label founder chromosomes, e.g.: founder “4” will have chromosomes labelled “4.1” and “4.2”. All founder chromosomes are labelled in this manner. The function iterates through all individuals. For every non-founder individual, we simulate meiosis in both parents and pass down a selected meiotic product from each parent. The notation inside the curly braces demonstrates how the haplotypes appear in the text file output. Segments are identified by their founder of origin (ID #), and the boundary positions are recorded in BP.

Other available gene-dropping software are not designed to efficiently simulate transmission of genomic regions through large genealogies. This is due to software either handling only few alleles (making them unable to simulate large regions), tracking a large number of alleles (making them inefficient for large genealogies and many replicates), or having additional functionalities (e.g., handling phenotypes). A detailed comparison to other software is provided in the Supplementary Appendix 4.

The output haplotypes describe the true IBD origin for each of the continuous segments that make up the simulated region. Hence, results obtained with our simulation function can help characterize patterns of genomic sharing between specific individuals and overall in the population. The output can be used to describe the distribution of haplotype lengths in the population or for specific individuals, obtain the distribution of the length of IBD segments shared by a pair of individuals, or estimate the likelihood of an IBD segment greater than a given length being transmitted from any ancestor. Simulation results can also be used to compare different statistical methods, as illustrated in Burkett, et al., 2022 who used a beta version of the function (with limited functionalities) to compare genomic- and genealogical/coalescent-based inference of homozygosity by descent in two different pedigree structures from the French-Canadian Founder population. In Supplementary Appendix 3, we provide additional examples of application of the function, highlighting all available functionalities.

## Conclusion

The gen.simuhaplo function combines the GENLIB R package’s existing support for handling large genealogies to allow users to simulate inheritance of large genomic regions even in genealogies with hundreds of thousands of individuals. To our knowledge no other simulators with similar functionalities supports such large genealogies.

