## Supplementary Material for "GENLIB: new function to simulate haplotype transmission in large complex genealogies"

### Appendix 1: Models for simulating meiosis

#### Model 1: Poisson process (Haldane, 1919)

If the user specifies “model=1” then a Poisson process will be used to simulate meiosis. The “params” argument of the function should be used to pass in a 2-element vector specifying the sex-specific recombination rates (in Morgans) of the Poisson process. For this model, the “params” argument should be the same as the “genetic_length” argument. This is because we use a Poisson process with rate L (length of region to be simulated in Morgans) to simulate the positions of the crossovers. Since a thinned Poisson process is also a Poisson process, we simulate the crossovers directly i.e., using a crossover process with rate L, instead of a chiasma process with rate 2L.

The number of crossovers is generated by sampling a Poisson distribution with rate L, the sampled number of crossovers are then uniformly randomly distributed over the length of the region (in genetic distance). The crossovers positions are then converted from genetic distance into physical distance. Once we have the physical locations of the crossovers, a meiotic product is created to pass down to the offspring.

#### Model 2: Zero-Truncated Poisson Count-Location model (Karlin and Liberman, 1978; Karlin and Liberman, 1979; Risch and Lange, 1979; Sturt, 1976)

If the user specifies “model=2” then a zero-truncated Poisson (ZTP) distribution will be used to generate the number of chiasmata:

$$p_{n}=\frac{e^{-\lambda}\lambda^{n}}{n!(1-e^{-\lambda})}, n>0$$

After obtaining the number of chiasmata, they will be distributed randomly along the simulated region, then the chiasma will be selected with p=0.5 to obtain the crossover positions. Note that this model guarantees a chiasma (obligate chiasma), but there still may be 0 observed crossovers.

The user must specify the parameter $\lambda$ of the ZTP distribution using the “params” argument. The user must pass in a 2-element vector to specify a parameter for each sex, which can be the same. One way to select an appropriate value of $\lambda$ would be to use the following equation and set E[X] to 2L (we use 2L since we are modelling chiasmata).

$$E[X]=\frac{{\lambda e}^{\lambda}}{e^{\lambda}-1}$$

To sample from this distribution, we sample a standard Poisson distribution with the same rate and resample if we obtain a value of 0.

#### Model 3: Stationary Gamma process (Broman and Weber, 2000)

If the user specifies “model=3” then a stationary Gamma renewal process as described in Broman and Weber (2000) will be used to simulate meiosis. The Gamma renewal process models the “interarrival” distances between chiasma using a Gamma(v, 2v) distribution, where v is the shape parameter, and 2v is the rate parameter. This restriction of 2v as the rate parameter keeps the expected value of the distribution to 1/2, for any average distance of 0.5 Morgans between chiasma. The user must specify the sex-specific values of v using the “params” argument.

The distance to the first chiasma is distributed differently than the rest of the inter-chiasma distances, which are distributed Gamma(v,2v). We obtain the distribution of the distance until the first chiasma (first arrival time) by selecting a distribution that will satisfy the stationary property. The stationary property means that the probability of chiasma formation will be equal anywhere along the chromosome, and the p and q terminals of the chromosome will be treated the same regardless which terminal we initiate the process at.

If the first arrival time is distributed according to the limiting distribution $\frac{\mathcal{F}(x)}{\mu}$of the interarrival distribution, then the renewal process will have the stationary property. Here $\mathcal{F}(x)$ is the survival function of Gamma(v,2v), and $\mu$ is the mean (1/2). This is equivalent to 2[1-F(x)] where F(x) is the cumulative distribution function (CDF) of Gamma(v,2v).

We evaluate the CDF using source code obtained from (https://people.math.sc.edu/Burkardt/cpp_src/asa239/asa239.html), which uses the algorithm for calculating the incomplete Gamma integral from (Shea, 1988). We generate 10,000 partial Riemann sums for 2[1-F(x)] for x values between 0 and the length of the region (in Morgans). Then we sample the 2[1-F(x)] using an inverse transform method using the partial sums.

After sampling the position of the first chiasma, we then sample the Gamma(v,2v) distribution (implemented in C++ std random library) to get the distance to the next chiasma. We continue this until the length of the region (in Morgans) is exceeded. We then independently select crossovers from the chiasmata, and convert the positions into BP, same as the other two models.

For all the models of meiosis, we use the C++ std random library for the Mersenne twister pseudorandom number generator, and the random distributions (Poisson, Uniform, Gamma).

#### Using a map to convert genetic distance into BP

Genetic distance does not perfectly correlate with physical distance along the simulated region. For instance, there are fewer observed crossovers around the centromere. In addition, the distribution of recombination sites seems to be different for males and females as well, e.g.: males have been observed to have higher rate of recombination around the telomeres (Bherer, et al., 2017). As discussed in Caballero et al. (2019), using maps to convert genetic distance into physical distance as part of simulating meiosis can alter the distributions of identity-by-descent (IBD) sharing between pairs of individuals of a given relatedness (Caballero, et al., 2019).

To specify a map to the gen.simuhaplo function, the map must be a data frame with a column named “cM” and a column named “BP”. Each row of the data frame should be an ordered pair that identify a point on the map, and the rows should be ordered from low to high (start to end of the simulated region).

When converting a crossover position from genetic distance to physical distance using the map, we find the pair of points that bound the crossover position, and use linear interpolation of the physical distances of the points to obtain the physical position of the crossover. This can be an expensive procedure if the map consists of many points (high resolution map) since we have to search this map for every meiosis. We recommend approximating the map with a few points as shown in Supplementary Figure 1.

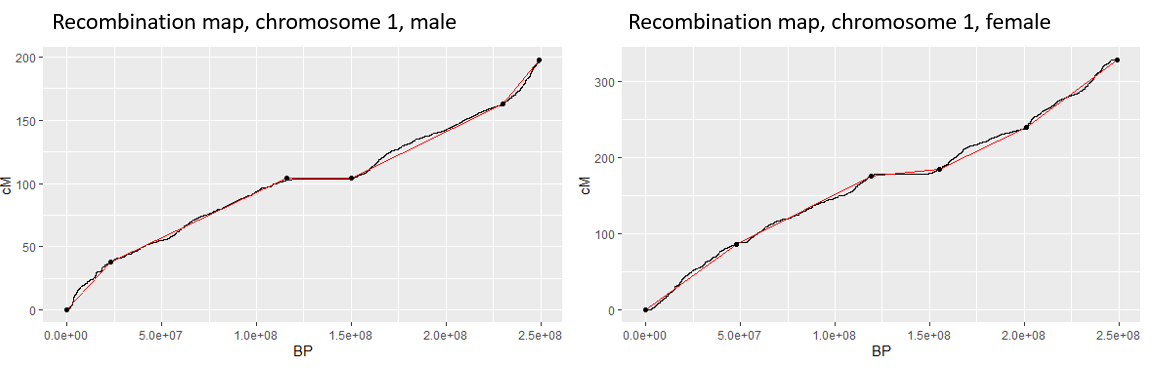

Supplementary Figure 1. High-resolution chromosomal map (black line) from Bherer et al., (2017). Maps are constructed from 100,000 + recombinations sampled from family studies. The red line shows the linear interpolation at a few specified points that can be more efficiently used by the simulation function.

### Appendix 2: Standalone scripts

The gen.simuhaplo function creates an output text file with the simulated haplotypes in the format shown in Figure 1 of the main text. We provide Python and Perl scripts that can be used to analyze the output of the function. The scripts can be found at: <https://github.com/R-GENLIB/simuhaplo_scripts>.

#### Tracing segments up the transmission path

The “IBD_traceback.py” script can be used for tracing back a segment in a proband’s simulated chromosome up the transmission path. The script requires two output files obtained from the gen.simuhaplo function: the “All_nodes_haplotypes.txt” output file (which is optionally created by gen. simuhaplo) and the “Proband_haplotypes.txt” output (automatically created by gen.simuhaplo). The script also requires a file containing the genealogy used for the simulations in standard pedigree format (i.e., individual id, father id, mother id, sex). This script simply identifies the boundaries of the segment to be traced back, and then follows the recombination history stored in “All_nodes_haplotypes.txt” upwards at each meiosis until reaching the founder.

In a theoretical, completely outbred pedigree there would only ever be one path connecting proband to founder. The inheritance of a segment from this founder would only depend on the number of meioses along that path, and the sex of the individuals along the path (if we consider sex-specific parameters of the meiotic process). One can analytically obtain a distribution for the lengths of inherited segments as described in (Boehnke, 1994) for a given path, however if there are multiple possible paths one would have to sum over all possible paths, and weight them according to the likelihood of inheritance down a given path.

Additionally, when deriving the distribution for the length of an inherited segment, it is considered that the meioses along the transmission path will potentially shorten the segment (Boehnke, 1994; Caballero, et al., 2019; Nelson, et al., 2018), but when crossing paths exist it is possible this assumption would be violated. If multiple paths converge at an internal ancestor, it is possible for the internal ancestor to be homozygous for the founder segment, and this ancestor may pass along a haplotype that has been lengthened by a recombination, instead of being shortened. This should be a very rare occurrence (depending on the levels of consanguinity), but the traceback script will be able to identify if it occurs.

The traceback script will let us check the exact path followed for the inheritance of a specific segment, in a given simulation. This information could be useful for identifying internal ancestors who play an important role in inheritance of mutations under study. The output file “All_nodes_haplotypes.txt” can get very large, and would not allow simulating millions of replicates in large genealogies. We provide the option of outputting the “All_nodes” files as a binary file to support millions of simulations in large genealogies. The standalone scripts currently use the text file output but we are working on writing equivalent scripts for the binary file, which will be made available at: <https://github.com/R-GENLIB/simuhaplo_scripts>.

#### Determining proportion of genome shared IBD

The “percent_IBD.py” script is used to compare the total proportion of the diploid chromosome that a pair of probands share IBD. The gen.simuhaplo function simulates both copies of a diploid chromosome for every individual, so when determining the percent shared IBD between a pair of individuals we must compare both copies to each other.

To do this, we extract the region from the output file, and create a list of all the continuous haplotype segments making up the simulated segment. Since we have a numerical identifier for the origin of each segment we then check if the two individuals share any of the same identifiers. If they do, we check the positions of these segments (which can be multiple discontinuous segments, or on different copies of the chromosome) and see if there is any overlap. We repeat this process for each of the shared identifiers. Once all overlapping regions are determined, we need to check if any of the identified IBD segments are directly adjacent to each other (i.e.: a longer IBD stretch inherited from a common internal ancestor, that is composed of multiple segments from multiple founders), then we consider it as one long segment.

When comparing the two individuals of a pair, both copies of the chromosome for individual 1 must be compared to both copies of the chromosome for individual 2. We do this by arbitrarily selecting one of the two individuals, ind1, and comparing the first copy of ind1 to both copies of ind2, the other individual in the pair. From this comparison we get a list of segments where ind 1/copy 1 overlaps ind2/copy1 or ind2/copy2. If any of the list of segments overlap (i.e.: ind2 was homozygous by descent (HBD)) then we merge the overlapping segments (not double counting, just taking the maximal boundaries of the region). We add up all the lengths of the segments to get total length that ind1/copy 1 is IBD to ind2/copy1 or ind2/copy2. We perform the same steps for ind1/copy2. The lengths are then added together to get IBD(ind1, ind2). We repeat the same steps after switching the place of the two individuals to get IBD(ind2, ind1). Finally, we average IBD(ind1, ind2) and IBD(ind2, ind1) to get the proportion shared IBD without double counting the HBD regions.

The “percent_IBD” script will identify the proportion shared between a pair, the number of discontinuous IBD stretches, the length of the longest stretch, and the ID’s for the founder(s) of origin for the IBD regions.

#### Converting simulated haplotypes into genotype data

The gen.simuhaplo function does not handle any genotype data, it only tracks the transmitted haplotypes with respect to their position and founder of origin (numerical identifier). We can convert this representation into genotype data using the “reconstruct.pl” script, given the haploid genotypes of the founders. We simply use the two haploid genotype data for the respective founder chromosomes to replace all the segments with their appropriate stretch of sequence data (using the BP positions of the segment). The haploid genotype information should be passed as a file with each line specifying a single founder chromosome. The file should be formatted such that each line starts with the numerical identifier for the founder chromosome, followed by a space, and then a string specifying the genotype. Haploid genotypes may be described as a string of any characters, so long as each founder chromosome has a string of the same length. Each single character in the string represents the genotype at a specific BP position. A second file should be provided that contains the BP position of each character (variant genotyped); the file should have the same number of lines as there are characters in the haploid genotype string, with only a single integer on each line (BP position).

### Appendix 3: Example Applications

Here we use our function to simulate haplotypes for probands in two genealogies. We perform 1000 replicate simulations and illustrate some of the analyses that can be performed with the simulation results and the additional scripts. We explore two genealogical structures of over 15 generations from the French Canadian founder population, both constructed using the BALSAC database (BALSAC project, Université du Québec à Chicoutimi, https://balsac.uqac.ca/) (Vézina and Bournival, 2020). The first genealogy was constructed from a sample of individuals recruited in a family study of asthma in the Saguenay-Lac-Saint-Jean (SLSJ) region (Laprise, 2014), which harbors higher levels of inbreeding due to its founding history and isolation. The second genealogy was constructed from a more out-bred sample from patients in ophthalmology clinics of Maisonneuve-Rosemont Hospital in Montreal (Varin, et al., 2017; Varin, et al., 2020). More details on the genealogies used in the example applications can be found in (Burkett, et al., 2022). Subsets of the present-day individuals were selected from the two genealogies to include in our simulations, yielding the genealogies shown in Supplementary Table 1. We simulate the equivalent of chromosome 1, with a length of 290,000,000 BP and genetic length of 198cM in males, and 328 cM in females.

Supplementary Table 1. Overview of the two genealogies used for simulations

|  | SLSJ | Montreal |
| --- | --- | --- |
| Number of probands included | 226 | 227 |
| Number of founders | 7608 | 9095 |
| Number of individuals | 55750 | 58942 |

#### Comparing the models of meiosis

To compare how the models of meiosis can affect the haplotype inheritance among probands we run 1000 simulations for each model of meiosis, both with and without the use of genetic-physical map. For the Poisson model we use 1.98, and 3.28 as our model parameters (length of chromosome in Morgans), for the ZTP model the parameters are 3.88, and 6.55, which we obtain from the equation in Appendix 1 to keep the same expected number of crossover, and for the Gamma model we use 4.5 and 5.5 as our parameters, which were the values found by (Broman and Weber, 2000) after fitting the Gamma model to their observed recombination data. The gamma model will always have expected number of crossovers equal to the Morgan length, so all 3 models have same expected number of recombinations per meiosis. The maps used are the approximations shown in Supplementary Figure 1.

Run times for the different meiosis models are shown in Supplementary Table 2. Using a more complicated Gamma model with a map adds ~1 minute to the simulation time compared to a simple Poisson model with no map.

Supplementary Table 2: run time, and output file size for 1000 simulations in both genealogies. Simulation was run on a i3-4160 CPU @ 3.60GHz machine. The function is currently only single thread.

|  | SLSJ Genealogy | | Montreal Genealogy | |
| --- | --- | --- | --- | --- |
|  | Time (min) | Output file size (Mb) | Time (min) | Output file size (Mb) |
| Poisson no map | 2.96 | 232 | 2.48 | 201 |
| Poisson w/ map | 3.08 | 230 | 2.70 | 200 |
| ZTP no map | 3.03 | 231 | 3.00 | 201 |
| ZTP w/ map | 3.23 | 230 | 3.02 | 200 |
| Gamma no map | 3.63 | 231 | 3.46 | 201 |
| Gamma w/ map | 3.75 | 230 | 3.56 | 200 |

We first compare the distributions of longest inherited founder segments between the different models of meiosis. For each of the 1000 simulation replicates, we obtain the length of the longest inherited founder segment for each proband over both copies of chromosome from all founders, yielding 227,000 segments for the Montreal genealogy and 226,000 segments for the SLSJ genealogy. The distributions are shown in Supplementary Figure 2. The ZTP and Poisson have almost identical distribution, which is to be expected since we are simulating a long chromosome where the chance of sampling 0 chiasma is low. The models that use the map have lower mean and more concentrated distribution. One explanation for this might be because the map sequesters crossovers into the tail regions of the chromosomes, and away from the centromere, this could lead to shorter segments after successive meioses. The gamma models also appear to have less dispersion and be more centered around the mean. This makes sense because the purpose of the gamma model is to account for chromosomal interference, i.e.: to space out recombination events, which would reduce the variance in segment length.

Next, we compare how the different models of meiosis may affect the proportion of IBD sharing. We check every pair of probands (25425 for SLSJ, 25651 for Montreal) to determine whether they share any part of the simulation segment IBD, i.e., we treat sharing as a binary variable. We calculate the frequency of IBD sharing as the proportion of the 1000 simulations where the pair had non-zero levels of IBD sharing. The distributions over all pairs are shown in Supplementary Figure 3. All models have very similar distributions, with the Gamma models showing slightly higher means (horizontal line).

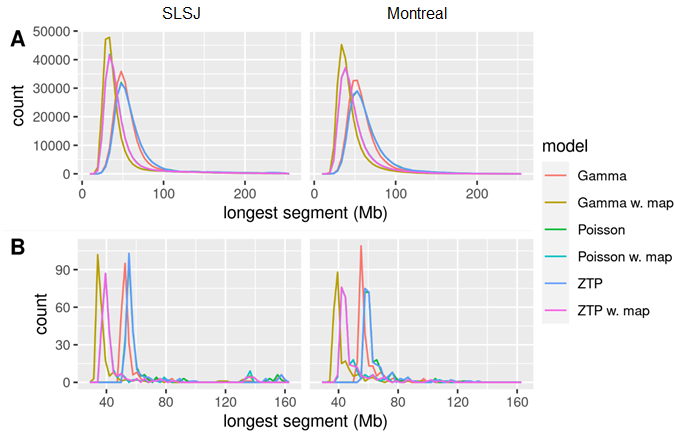
Supplementary Figure 2. A: length of longest segment for all probands, all simulations. B: longest segment, average value for each proband (averaged over simulations).

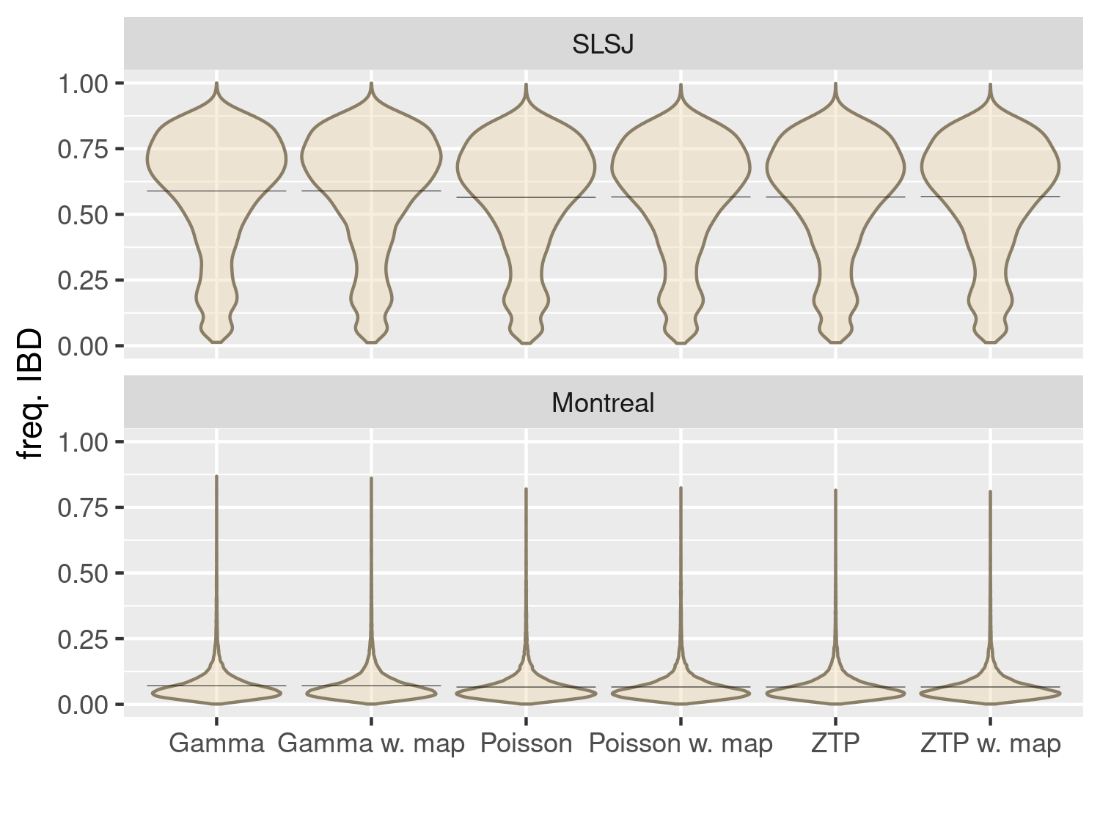
Supplementary Figure 3. Distribution over all pairs (25425 for SLSJ, 25651 for Montreal) of the frequency of IBD sharing (proportion of the 1000 simulations where the pair had non-zero levels of IBD sharing) for the different meiosis models.

#### Exploring IBD sharing between pairs of probands

In this example application, we only consider the simulation results from the Poisson model with no map. We check each pair of probands using the “IBD_percent.py” script and obtain details on the IBD sharing between each pair of probands for each of the 1000 simulations. We also obtain a genealogical kinship coefficient between every pair of individuals using the gen.phi command in GENLIB. By definition, the mean of the proportion of the genome shared IBD between a pair should be equal to twice the kinship. We obtain the proportion of the simulated segment shared IBD for each of the 1000 simulations, and compare the mean to the kinship in Supplementary Figures 4 and 5.

As expected, the mean proportion of the simulated segments shared IBD correlates strongly with kinship, with slope 2 and intercept 0, and the variance increases as kinship increases. The patterns of variance can be seen in the zoomed plot (Supplementary Figure 5). In addition, especially in the Montreal genealogy, it seems that the variance is higher, at a given kinship level, for pairs with higher mean proportion of segments shared IBD. This pattern is less apparent in the overall more related SLSJ population.

For a given pair the kinship can only tell us the expected value of the proportion of the genome shared IBD. The distribution can be obtained from the simulation results. A specific value of the kinship coefficient can be realized in different ways, e.g.: a pair of individuals that share a grandparent, and a pair of individuals that share two great-grandparents will have the same kinship coefficient and therefore the same expected value of total percent shared IBD. However, the distributions of the genomic proportion shared IBD may be quite different. To explore this, we selected 4 pairs of probands in the SLSJ genealogy. Pair 1 has kinship similar to pair 2 and pair 3 has kinship similar to pair 4. The difference in the distribution of the simulated segments shared IBD is shown in Supplementary Figure 6. Both sets of pairs have similar kinship values, but different distribution of IBD sharing. Pair 4 and pair 2 realize their kinship earlier in the genealogical history, which explains their higher frequency of no IBD sharing, i.e., when sharing occurs, it will be a longer segment but there is a higher probability of sharing not occurring since it is largely due to a fewer lower generation ancestors, instead of spread out among more shared ancestors at a higher generation.

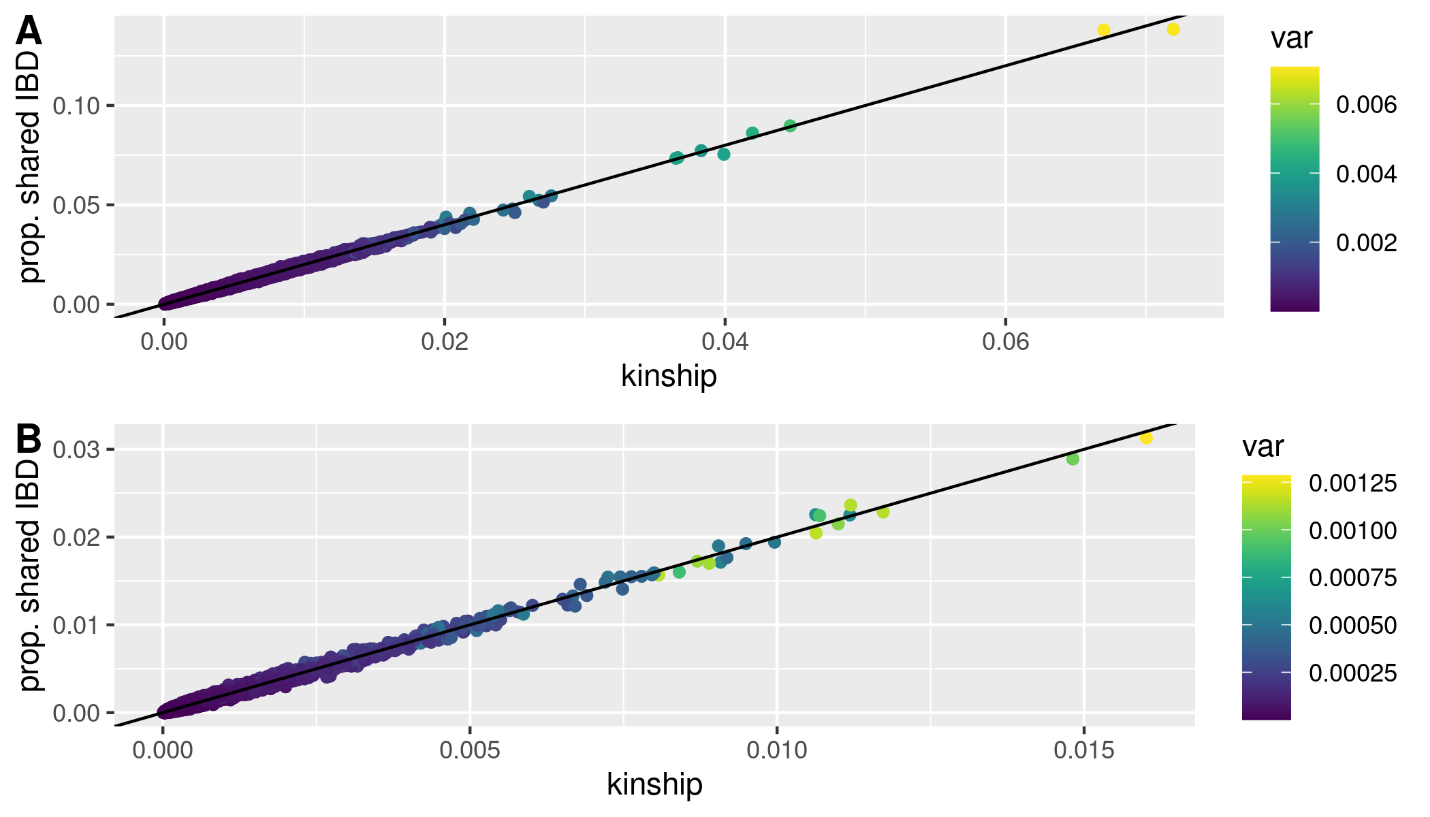

Supplementary Figure 4. Genealogical kinship vs. mean proportion of the simulated segment shared IBD. A: SLSJ genealogy, 25425 proband pairs (226 probands). B: Montreal genealogy, 25651 proband pairs (227 probands). The colours show the variance over the 1000 distribution (var). The expected regression line of slope=2 and intercept=0 is shown.

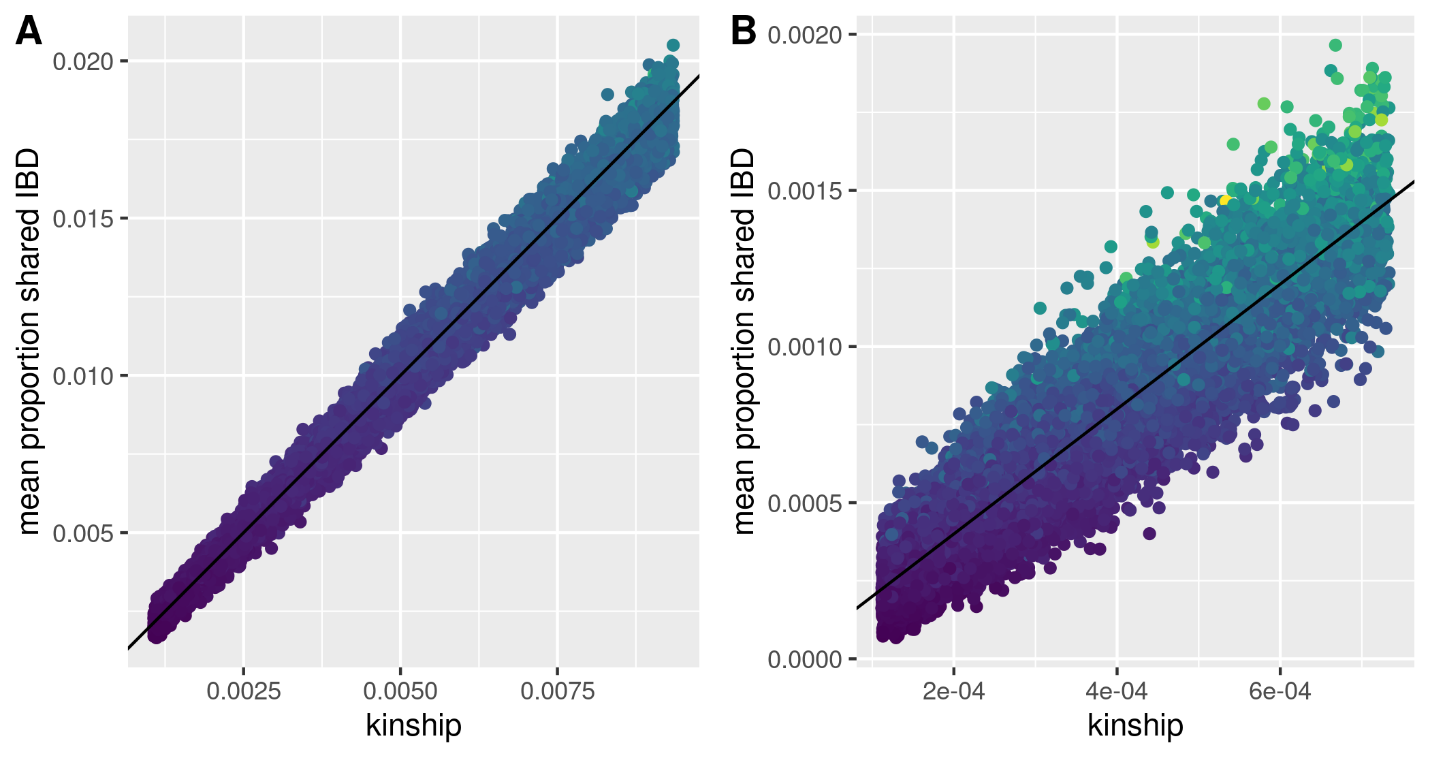

Supplementary Figure 5. Genealogical kinship vs. mean proportion of the simulated segment shared IBD, zoomed to remove the bottom and top 10 percentile of kinship values. A: SLSJ genealogy, 25425 proband pairs (226 probands). B: Montreal genealogy, 25651 proband pairs (227 probands). The colours show the variance over the 1000 distribution (var legend shown in Figure 4). The expected regression line of slope=2 and intercept=0 is shown.

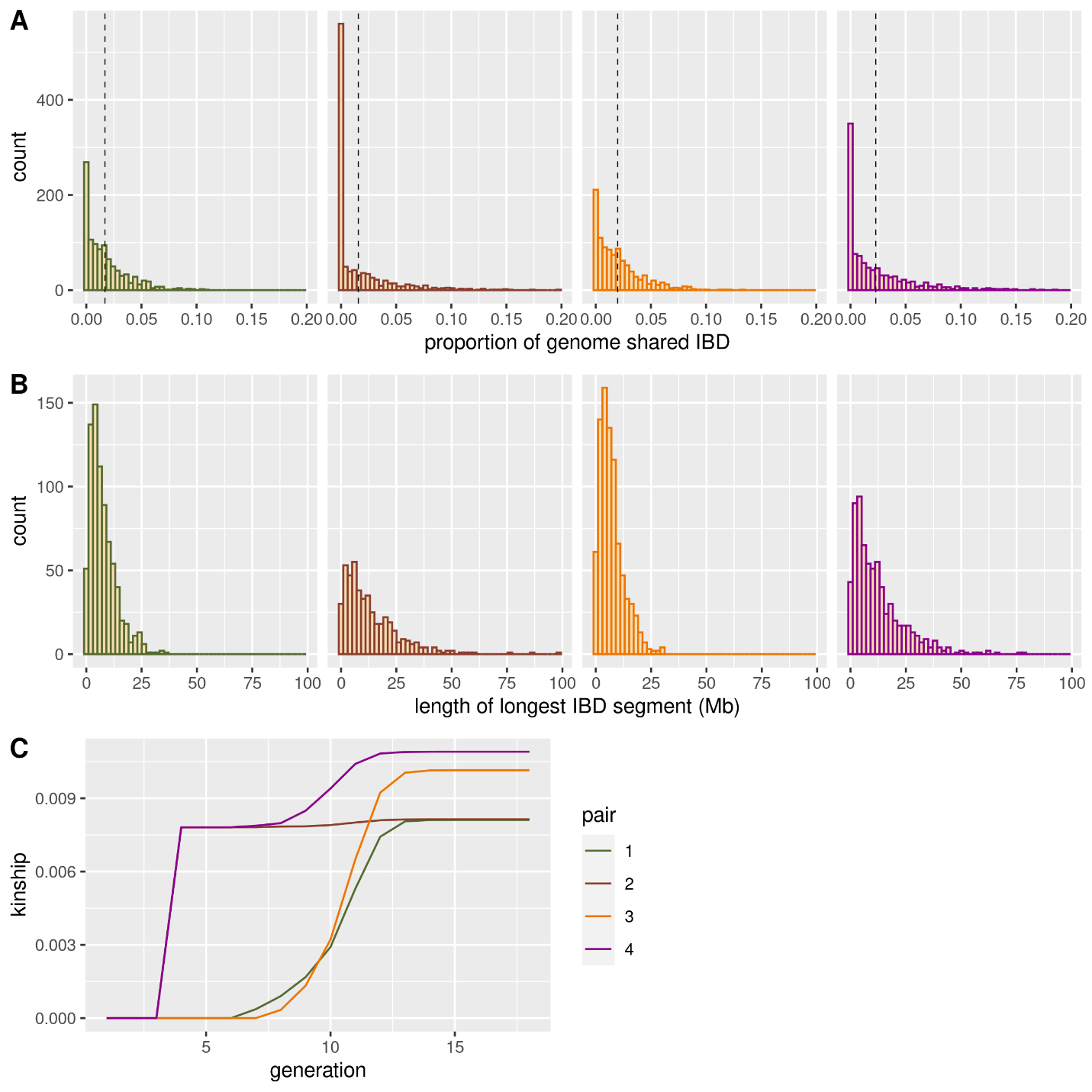

Supplementary Figure 6. IBD sharing for 4 arbitrarily selected pairs in the SLSJ genealogy. A: histogram of the proportion of the simulated segments shared IBD values over 1000 simulations (the mean is indicated by the dashed line). B: histogram of the length (in mega base pairs) of the longest IBD segment (for those simulations with non-zero sharing) C: Genealogical kinship by generation for the 4 pairs of probands.

#### Inheritance of IBD segment from ancestor

As mentioned earlier, it is difficult to analytically derive distributions for lengths of inherited IBD segments for a given proband-founder relationship in consanguineous populations. The simulation results will estimate this distribution, and using the traceback script we can further obtain a distribution for each possible path of inheritance.

We ran simulations on a small sub-tree of the Montreal genealogy, we only consider the connections between proband “222” and their ancestor “335” (Supplementary Figure 7). We ran 25,000 simulations (segments not originating from “335” are all given the same identifier of “0”). If “222” inherits a segment from “335” then we can use the traceback script to identify which transmission path was followed. We can also detect any instance of the concatenation event discussed in Appendix 2.

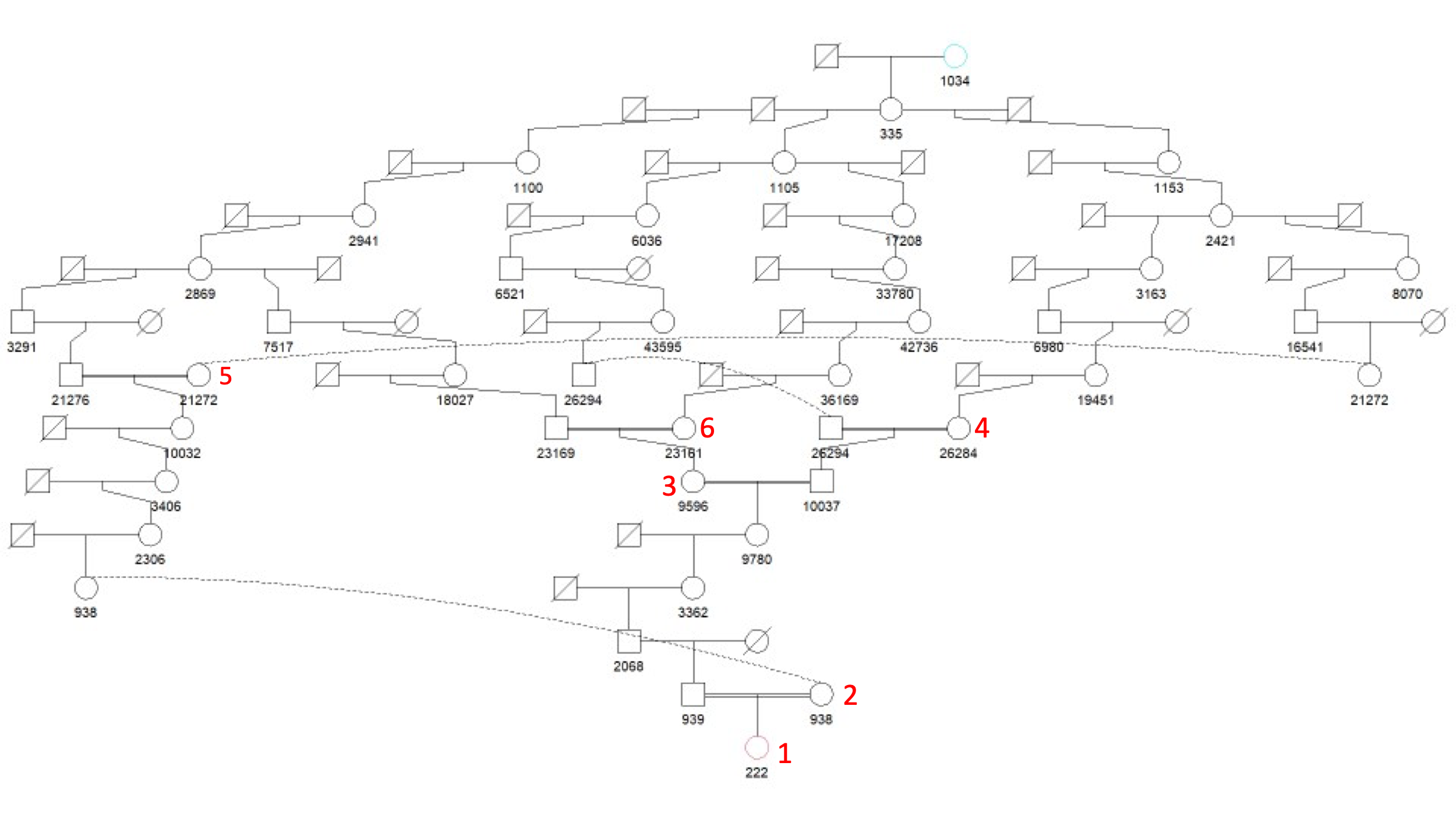

Supplementary Figure 7. Genealogy between proband “222” and ancestor “335”. There are 6 possible paths between “222” and “1034”, as labelled in the figure.

Of the 25,000 simulations, there were 4968 instances of a segment being inherited from ancestor “335”. Of those 4968 simulations, 4961 involved inheritance directly down a single path. The frequency of each path taken is shown in Supplementary Table 3. The remaining 7 occurrences were inherited through a ‘concatenation’ event, i.e.: multiple paths joined at an internal node. Path 2 and 5 are shorter, leading to their increased prevalence. Even though the ‘concatenation’ event was quite rare, when it occurs it leads to a much longer segment (Supplementary Figure 8).

Supplementary Table 3. Frequency of each possible paths between proband “222” and ancestor “335” described in Supplementary Figure 7.

| Path1 | Path 2 | Path 3 | Path 4 | Path 5 | Path 6 |
| --- | --- | --- | --- | --- | --- |
| 650 | 1396 | 403 | 405 | 1401 | 436 |

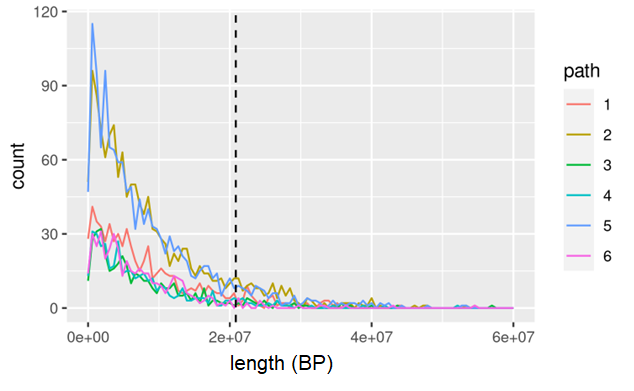

Supplementary Figure 8. Distribution of inherited segment lengths, depending on the path of inheritance between proband “222” and ancestor “335” described in Supplementary Figure 7. Horizontal line is the mean of the 7 segments that were inherited through concatenating segments at one of the internal ancestors.

### Appendix 4: Comparison to other software

Use of simulation tools is widespread throughout human and population genetics, animal and plant breeding. Simulation approaches can be considered as either backward-in-time, or forward-in-time. Forward simulators are used to simulate the progression of a population under some models of fitness, selection, mating, population size, mutation rates etc. Many tools do not let users specify a pedigree since they are meant to simulate the mating.

Most of the gene-dropping tools that follow user-specified pedigrees are created to track specific alleles, evaluate phenotypes, or condition on observed data (Li, et al., 2015; Nieuwoudt, et al., 2020). Since our tool only tracks position and origin of chromosomal segments it is more lightweight and can efficiently simulate very large, complex genealogies.

Libiger and Schork (2007) published a gene-dropping tool to investigate IBD segment sharing. Their approach tracks states of particular marker alleles as opposed to the positions of segments. We could not locate software corresponding to the paper. SimPed (Leal et al., 2005) is another such software that performs gene-dropping on individual marker loci. The software was obtained from (<https://www.hgsc.bcm.edu/software/simped>) and we attempted to use it to simulate 25 hypothetical markers, each spaced 0.1 Morgan apart for our Montreal genealogy. The software had a hard-coded maximum of 3000 individuals per genealogy. After altering this and recompiling the source code, the program crashed while attempting to simulate results for our genealogy. On small sample genealogies with the same parameters the software worked as intended.

XSim, as described in Cheng (2015) uses the approach of tracking positions and origins of chromosomal segments, however the software (<https://github.com/reworkhow/XSim.jl>) is designed for simulating breeding schemes, and mating programs, and not for dropping down user defined pedigrees.

The available software most similar to ours are PedSIM (Caballero, 2019) and IBDsim (<https://cran.r-project.org/web/packages/IBDsim/index.html>). However, both tools are designed for the evaluation of close relatives (small pedigrees).

PedSim has a unique file format specification for the input pedigrees. The file format as described at (<https://github.com/williamslab/ped-sim#def-fileigree simulator>) is not amenable for genealogies with consanguineous loops, or individuals that appear in multiple generations. The script provided in the github repository to convert PLINK “*.fam” file format to the PedSim file format fails to convert our example genealogy.

IBDsim, which is part of the Pedsuite (<https://cran.r-project.org/web/packages/pedsuite/index.html>) set of tools crashes when using the ped() command to try to load in our example pedigrees.
